# Spatially segregated responses to visuo-tactile stimuli in mouse neocortex during active sensation

**DOI:** 10.1101/199364

**Authors:** J Couto, S Kandler, D Mao, BL McNaughton, L Arckens, V Bonin

## Abstract

Multisensory integration is key for perception and animal survival yet how information from separate senses is integrated has been debated for decades. In the cortex, information from each sense is first processed in primary sensory areas and then combined in association areas. An alternative hypothesis to this hierarchical model is that primary sensory cortices partake in multisensory encoding. We probed tactile and visual responses in primary somatosensory and visual cortices in awake behaving animals using two-photon calcium imaging from layer 2/3 excitatory neurons. In support of an hierarchical model we found segregation of visual and tactile responses. Tactile stimuli evoked responses in S1 neurons. In striking contrast, V1 neurons failed to respond to tactile stimuli. This was true for passive whisker stimulation and for stimulation during active whisking. Furthermore, responses of V1 neurons to congruent visuo-tactile cues during active exploration, a condition where vision precedes touch, were completely abolished in darkness. The rostro-lateral area of the visual cortex responded to both visual and tactile aspects of the stimuli and may form a substrate for encoding multisensory signals during active exploration. Our results indicate that primary sensory areas mainly encode their primary sense and that the impact of other modalities may be restricted to modulatory effects.

## INTRODUCTION

Animal behaviors rely on information from multiple senses. Multisensory integration enhances processing of weak or ambiguous sensory stimuli and supports sensory perception and learning (Gingras et al., 2009; Olcese et al., 2013; Stein and Stanford, 2008). For example, visuo-tactile congruence improves discrimination performance (Pasalar et al., 2010) and visual motion impacts tactile motion perception (Bensmaïa et al., 2006). In cortex, encoding of sensory information is thought to occur in primary sensory areas that are dedicated to processing independent senses and then combined in cortical association areas (Andersen et al., 1997; Bruce et al., 1981). An alternative hypothesis is that primary sensory cortices encode information from multiple senses (Ghazanfar and Schroeder, 2006; Kandler, 2015; Schroeder and Foxe, 2005; Stein and Stanford, 2008). Yet, empirical evidence that neurons in primary sensory areas respond to inputs from their non-preferred sensory modality is scarce.

Electrophysiological and functional imaging studies claim that the primary visual cortex is subject to multi-sensory influences (Murray et al., 2016; Zangaladze et al., 1999). Recognition of tactile objects evokes activity in primary visual cortex of both blind and sighted subjects (Amedi et al., 2010; Merabet and Pascual-Leone, 2010; Sathian, 2005) suggesting a role in normal sensory function and in recovery from sensory loss.

The rodent primary visual cortex (V1) receives projections from other sensory areas (Henschke et al., 2015; Iurilli et al., 2012; Massé et al., 2016; Van Brussel et al., 2011) and multi-modal stimulation impacts neural activity in V1 (Bieler et al., 2017b; Iurilli et al., 2012; Kayser et al., 2008; Vélez-Fort et al., 2018; Wallace et al., 2004). Moreover, dedicated neuronal populations in V1 have been reported as targets of auditory inputs (Ibrahim et al., 2016) and pyramidal neurons in S1 and A1 target vasoactive intestinal peptide neurons in V1 (Fu et al., 2014). In addition to encoding visual inputs, activity in the primary visual cortex of the mouse reflects arousal state (Vinck et al., 2015) and locomotion (Niell and Stryker, 2010). Mice use both vision and touch in virtual reality navigation behaviors (Sofroniew et al., 2014), yet it is not known how these senses are combined during exploratory behaviors. Moreover the tactile thalamus but not the visual thalamus of anesthetized rats is capable of multisensory integration suggesting that multisensory integration may occur at early sensory processing stages (Bieler et al., 2018).

We investigated the impact of visuo-tactile stimuli in mouse primary somatosensory (S1) and visual cortex (V1) during voluntary locomotion. Passive stimulation of the whiskers suppresses V1 neural activity (Iurilli et al., 2012), however activity increases have been reported during tactile discrimination and object exploration (Vasconcelos et al., 2011). Whether tactile inputs alone drive activity in V1 has been a matter of debate (Bieler et al., 2017a; Vasconcelos et al., 2011; Wallace et al., 2004).

Here, we aimed to probe the extent to which neurons in V1 respond to tactile stimuli by recording activity from thousands of neurons in layer 2/3 to visual and tactile sensory stimuli during voluntary locomotion. We devised a locomotion assay to probe responses to congruent visuo-tactile stimuli during active sensation. We found that neurons in V1 are essentially activated by stimuli of their primary sensory modality. This was the case even during active exploration of visuo-tactile cues. These results highlight the unimodal nature of V1 sensory responses and imply that stimuli of other sensory modalities may be restricted to modulatory influences on cellular activity.

## RESULTS

### Primary visual cortex responds to visual but not whisker stimuli

To investigate the impact of tactile stimuli in the primary visual cortex during passive stimulation we performed calcium imaging in awake, head-fixed mice. After habituation to head fixation, mice underwent experiments for mapping visual and tactile evoked responses. Visual stimulation was delivered through a screen, spanning 0-120 degrees of the contralateral visual field. The contralateral whiskers were stimulated with air-puffs (Figure 1A). To reveal the spatial extent of activation by stimuli from the distinct modalities and map the location of the visual and somatosensory cortices, we performed widefield calcium imaging on transgenic mice that express the calcium indicator GCaMP6s in cortical excitatory neurons (Thy1-GCaMP6s or TRE-GCaMP6s-CaMKII). Visual stimulation for retinotopic mapping consisted of moving bars or a circular patch that drifted across the screen. Visual stimuli evoked activity in the visual cortex and air-puff stimuli activated the somatosensory cortex (Figure 1B) suggesting little spatial overlap in the cortical regions activated by visual and tactile stimuli.

**Figure 1:**
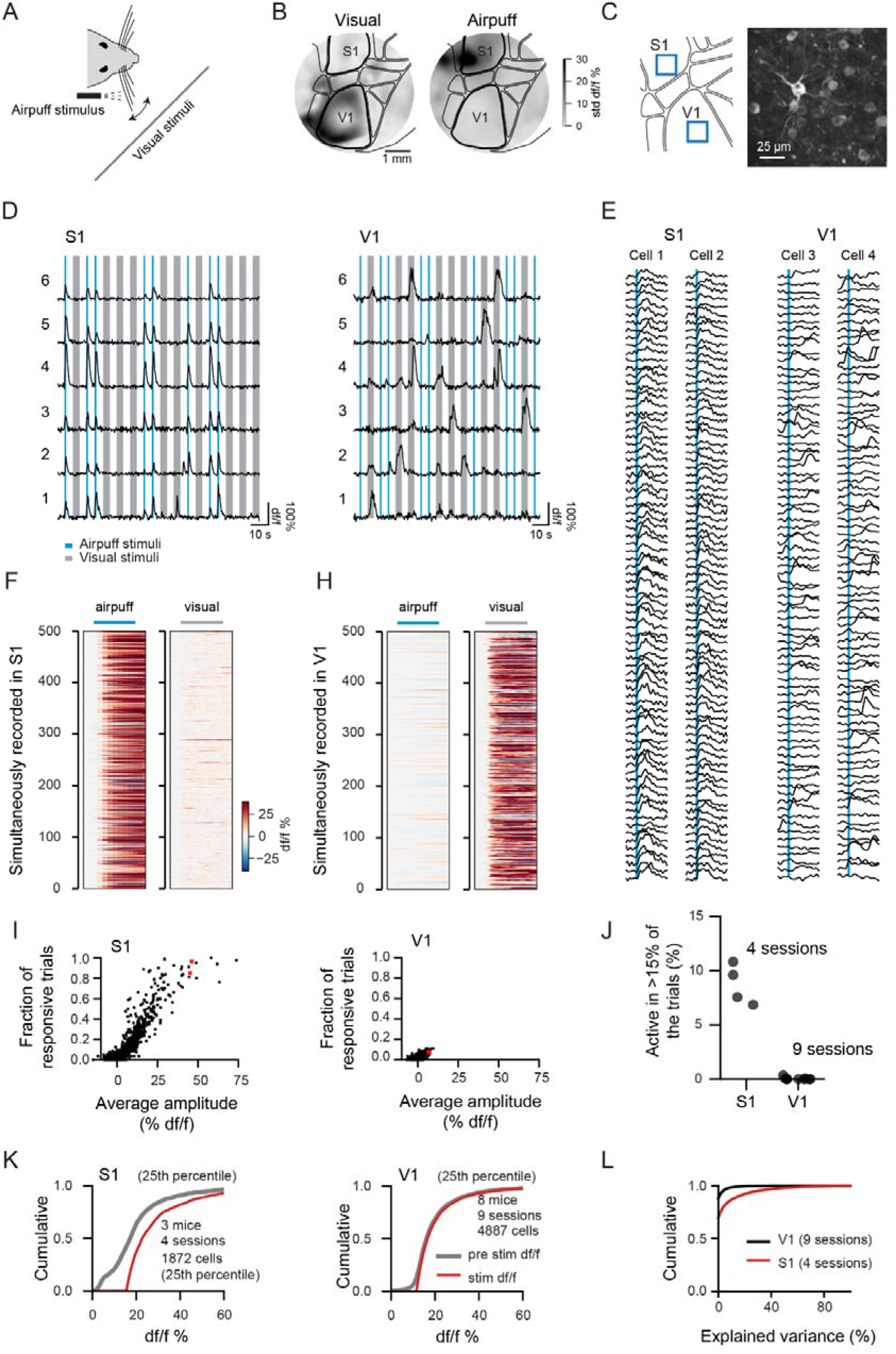
Primary visual cortex does not respond to passive whisker stimulation. A – Illustration of the experimental setup for whisker and visual stimulation. B – Wide-field imaging through a chronic cranial window. Color code is the standard deviation of the normalized fluorescence change during the stimulation period (8s for visual - 20 deg patch circling the screen edges) and 1s for whisker air puff stimulation). Left is for visual and right for air-puff stimulation. C – Illustration of sampled targeted locations for two-photon cellular calcium imaging and max projection of a recording in S1. D – Example of raw traces from individual neurons responding to the air puff stimulus in S1 (right) and to the visual stimuli in V1 (left). Blue and grey shaded areas are the air puff and visual stimuli. E – Responses of individual neurons to multiple trials of the air puff stimulus in somatosensory cortex and in primary visual cortex (not locked to the stimulus). F – Average responses of the 500 cells with highest amplitude (stim-pre df/f) to the air puff stimulus for a recording session in somatosensory cortex. The baseline before the stimulus was subtracted to highlight potential responses. Left – response to the air puff. Right – response to the visual stimuli (same cells as in the left). H – Same as F but for primary visual cortex. I – fraction of responsive trials versus the average df/f amplitude during the air puff stimulus for cells in the somatosensory(right) and primary visual cortex (left). Red dots are the cells in E. J – Percentage of cells responsive during 15% of the trials for individual recording sessions. K – Distributions of df/f amplitudes before (grey) and during (red) the air puff stimulus for cells in the 25th upper percentile of df/f amplitude. L – Distributions of explained variance by the average response to the air puff stimulus for the sessions in K.

We then mapped responses to passive stimulation at single neuron resolution using two-photon calcium imaging (Figure 1C). To probe the responsiveness to visual stimuli, we used full-field bandpass noise with different combinations of spatial and temporal frequency. Tactile stimuli consisted of trains of air puffs lasting 1 second (20ms pulses at 10Hz). We recorded from 7486 neurons in somatosensory cortex (4 sessions in 3 mice) and from 20595 cells in primary visual cortex (9 sessions in 8 mice). Neurons in the somatosensory cortex responded to air puff stimulation of the whiskers but not to the visual stimuli (Figure 1D – 500 most responsive to the air puff stimulus taken from one experiment). In contrast, neurons in visual cortex were not activated by whisker stimulation despite often being responsive to visual stimulation (Figure 1D - right). We used 40-80 trials to assess whether neurons were responsive to the air puff stimulus (Figure 1E). We considered responsive trials as those were the activity during the air puffs was larger than twice the standard deviation of the activity preceding the stimulus. Neurons in somatosensory cortex but not in primary visual cortex responded to repeated presentations of the air puff stimulus (Figure 1I). While in S1, 621 out of 7486 cells were active in at least 15% of the air puff stimulus trials, in V1 only 9 out of 20595 cells where active under the same criteria (Figure 1J). The absence of responses to the air puff stimuli was not related to biased sampling of the cortical space since multiple locations in V1 were sampled in the same mouse (Supplementary Figure 1A).

To compare the effect of stimulation on cellular activity across all experiments, we selected cells in the 25^th^ upper percentile of amplitude, calculated using the average df/f during the stimulus. While in somatosensory cortex the distributions of average df/f preceding and during the stimulus epoch were significantly different (Kolmogorov–Smirnov test, p-value < 0.01), this was not the case for cells recorded in the visual cortex (Figure 1K, see Supplementary Figure 1B for a comparison of all cells). Similar results were obtained when comparing explained variance by the average response to the air puff stimuli (Figure 1L). These results suggest that there is no cellular activation by passive whisker stimulation in layer 2/3 excitatory neurons in the primary visual cortex. Tactile activation of cells in primary visual cortex could require active whisking. We therefore devised an assay for probing whisker responses during active whisking.

### Probing cortical responses to congruent visual and tactile stimuli

To test whether cellular tactile activation occurs for ethologically relevant stimuli, we devised an experimental assay where (1) visual and tactile stimuli are congruent and (2) sensation is gated by animal locomotion. Somatosensory responses in S1 during active whisking are stronger and longer lasting than during passive stimulation (Krupa et al., 2004). Sets of cues were attached to the treadmill belt at discrete locations and were visible 20 cm before contacting the whiskers (Figure 2A). During locomotion, mice extended their whiskers which contacted the cues (Figure 2B). For comparison with the air puff stimulus, a short duration (1s) air puff was delivered at a fixed location of the belt in these experiments as well. Animals were trained for head-fixed locomotion, for a period of at least 2 weeks, after which they often performed over 100 laps/hour at an average speed in locomotion bouts of 40 cm/s (Figure 2C).

**Figure 2:**
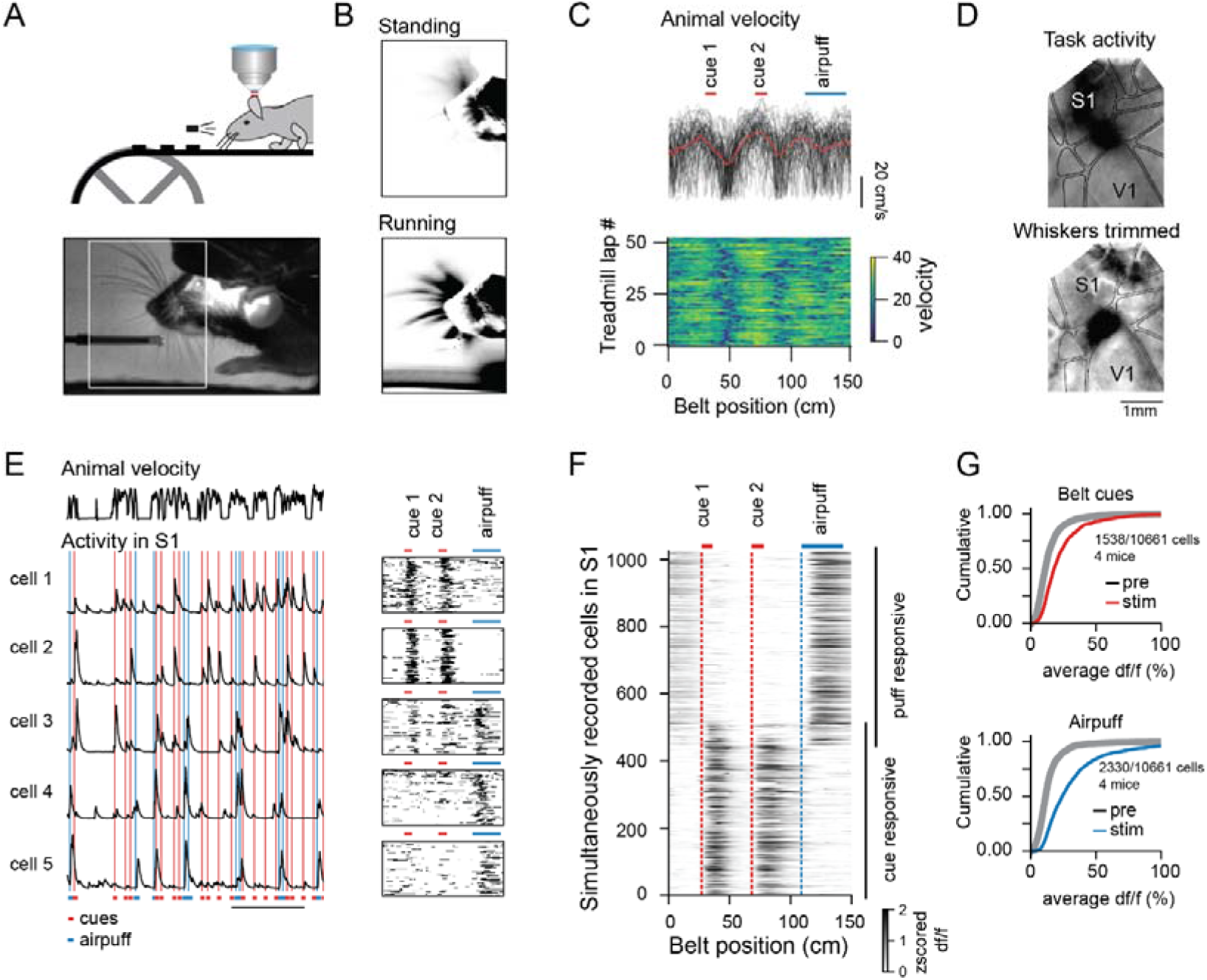
Somatosensory cortex responds to visuo-tactile cues during active whisking and locomotion. A – Illustration of the experimental setup and camera frame during locomotion. B – Whisker motion during locomotion. Top – Average difference between frames for the region in A during standing bouts. Bottom – Same as top for locomotion bouts. C – Animal velocity in locomotion bouts. Top – Average locomotion during an experiment (red) and individual laps (black). Bottom – Animal velocity is stereotypical across laps. D – Activity in somatosensory cortex is largely driven by the whiskers. Top – Standard deviation of wide-field calcium fluorescence during locomotion on a cued belt. Bottom – Same as top but after trimming the whiskers. E – Example of cellular activity in somatosensory cortex. Left – Raw calcium traces of 5 cells. Right – Deconvolved activity maps for different laps. Location of the cues (red) and the air puff (blue) stimuli. F – Average z-scored activity from responsive cells from one experimental session. G – Distribution of cue (top) and air puff (bottom) amplitudes for responsive cells in recorded in somatosensory cortex.

To measure the spatial extent of cortical activation by visuo-tactile cues we performed widefield calcium imaging. Mapping calcium activity during locomotion (animal velocity > 1cm/s) to the position on the treadmill revealed activation of the primary somatosensory cortex and anterior visual cortex (Figure 2D top, supplementary movie 1) at cue locations. To isolate the sensory source of activation we trimmed the whiskers (1 mouse). Activity in the somatosensory cortex at cue locations was abolished by whisker trimming suggesting a strong whisker-driven component of cortical activation. Nonetheless, anterior regions of the visual cortex remained active even after whiskers were trimmed (Figure 2D bottom).

To investigate cellular responses during active sensation, we performed two-photon calcium imaging. The goal of these experiments was to establish a baseline in somatosensory cortex to compare with cellular activity in visual cortex. Responses to the cues in somatosensory cortex evoked calcium transients repeatedly in individual trials (Figure 2E). In individual experiments, a large fraction of cells was driven by the cues or the air puff stimuli and a smaller fraction by both (Figure 2F). The average amplitude of calcium transients before and after the stimuli was significantly different (Kolmogorov–Smirnov test, p-value < 0.001). Across 4 mice, 1538 out of 10661 responded significantly to the cues and 2330 out of 10661 to the air puff stimuli (KS test, p<0.01) in S1.

Thus, the assay drives strong activity in somatosensory cortex. We then investigated responses to treadmill cues in the visual cortex.

### Anterior areas of the visual cortex respond to the visual aspect of the cues

We hypothesized that, if present, somatosensory responses would be strongest in anterior areas of the visual cortex, i.e. closer to the primary somatosensory cortex. Therefore, we recorded cellular activity using two-photon cellular calcium imaging from the anterior visual cortex in the border between primary visual cortex and the rostro-lateral area. We found 143 out of 3220 excitatory layer 2/3 neurons (in an individual session) responding to the cues (Figure 3A,C). We then recorded from the same neurons in complete darkness (lights turned off without time for light adaptation). All responses vanished in the darkness condition suggesting that cells respond to the visual aspect of the cues (Figure 3B,D). These results suggest that the activity we reported in anterior areas of the visual cortex after whisker trimming has a visual origin (Figure 2D – bottom). In addition, widefield activity in light and darkness conditions activated different locations of anterior visual cortical areas (Supplementary Figure 3A). Scotopic light conditions strongly suppressed responses to visuo-tactile cues and the distribution of average fluorescence amplitude triggered to cue position in scotopic conditions was indistinguishable from that measured before the cues.

**Figure 3:**
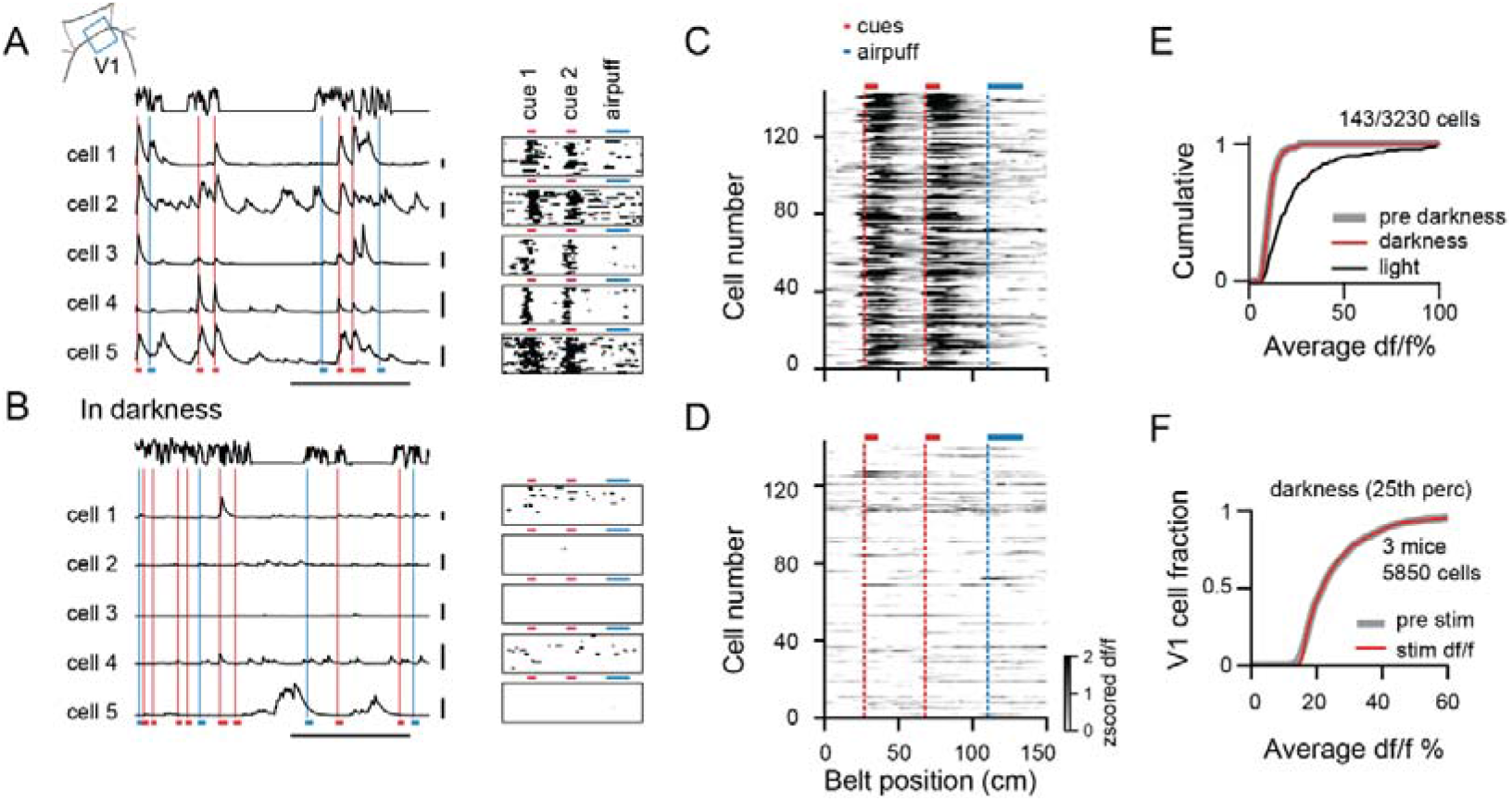
Responses to visuo-tactile cues in visual cortex vanish in darkness. A – Responses to visuo-tactile cues of 5 example cells. Left – raw df/f. Right – deconvolved activity across laps. B – Responses of the same cells as in A in scotopic conditions. C – Average response to the belt of 140 cells responding to the cues. D – Response of the same cells as in C in darkness. E – Quantification of average df/f for cells responding to visuo-tactile cues. F – Responses of cells in the 25th percentile in the darkness condition. Gray – average before the cue. Red – average during the cue.

We then asked whether tactile signals drive visual cortex activity during active whisking when the cues are not in visual reach. We therefore conducted experiments with a visual shield located 2 cm in front of the whiskers that blocked the view to the treadmill while allowing tactile exploration. In this condition, we recorded from the primary visual cortex from 3 mice (5850 cells) and found no evidence for somatosensory activation (Figure 2B). These results could be specific to the classes of excitatory neurons labelled in the Thy1-GCaMP6s and TRE-GCaMP6s-CaMKII mouse lines. We therefore injected an adeno-associated virus carrying GCaMP6m under the human synapsin promotor in visual cortex (3 mice with injections in V1) and found no significant difference between the distributions of average df/f in the time preceding and during cue crossing (Supplementary Figure 3D; 25.5±0.3 before; 25.8±0.3 during; mean±s.e.m.; Kolmogorov–Smirnov test p-value 0.18).

These results suggest that V1 layer 2/3 neurons do not encode cross-modal signals related to active whisking.

## DISCUSSION

We used a head-fixed treadmill assay with controlled tactile and visual stimuli to probe tactile signals in mouse primary visual and somatosensory cortices during voluntary locomotor behavior. We observed activation of layer 2/3 neurons in cortical areas V1 and S1 exclusively by stimuli of their primary sensory modality (Figure 1). Tactile activation was limited to neurons in S1 and higher order visual areas, observed both in darkness and under photopic conditions and disrupted by whisker trimming (Figure 2 and 3). V1 neurons were activated by visual features of the treadmill cues since responses were only observed under photopic conditions (Figure 3). Control experiments in darkness, showed that V1 and S1 neurons do not encode signals from their non-preferred modalities. These results show that cellular responses in primary sensory areas are essentially unimodal, suggesting that multisensory integration may occur in higher sensory and/or associative areas.

Cellular evidence of cross-modal activity in the rodent visual cortex stems primarily from electrical recordings of neuronal responses to passive stimulation (Bieler et al., 2017a, 2017b; Ibrahim et al., 2016; Iurilli et al., 2012; Wallace et al., 2004). A study of auditory stimulation (Ibrahim et al., 2016) reported activation of layer 1 inhibitory neurons but only modulations in layer 2/3 neurons. These studies used few stimuli and could not address the degree to which V1 neurons encode diverse information about distinct sensory modalities. Defying previous reports claiming that ~4% of neurons in primary visual cortex respond to somatosensory stimuli (Wallace et al., 2004), in awake mice we could not evoke tactile responses in primary visual cortex. Differences may lie in the different experimental assays used as previous evidence stems from electrical recordings in anaesthetized rats and stimuli delivered mostly though skin touch. The two-photon calcium imaging approach that we employed in behaving mice has a higher yield and sensitivity, allowing us to probe responses of large populations of neurons simultaneously with high spatial resolution and without potential confounds induced by anesthesia.

One study in freely moving rats (Vasconcelos et al., 2011) showed, in darkness, V1 activity linked to animal location and to objects as well as modulations correlated with tactile discrimination behavior. In our experiments, we found responses in darkness, only when enough time had passed for the animal’s eye to adapt to the infrared illumination (735nm). This is evidence of the high sensitivity of mouse vision and may raise concerns about studies that claim activity in darkness without controlling for light adaptation.

Long-range cortical inputs from somatosensory cortex to visual cortex have been documented (Charbonneau et al., 2012; Kim et al., 2015; Masse et al., 2016; Stehberg et al., 2014; Van Brussel et al., 2011) yet we did not find evidence for responses to whisker air puffs or tactile cue onsets in primary visual cortex. There are two main potential explanations for this: 1) somatosensory projections to primary visual cortex may carry non-sensory information or 2) excitatory neurons in layer 2/3 of the primary visual cortex do not receive direct and or only weak inputs from the somatosensory cortex. In addition, anterior visual cortical areas A and RL are interconnected with both the somatosensory barrel cortex and the primary visual cortex (Wang et al., 2012) and respond to both visual and tactile stimulation (Olcese et al., 2013). Neurons in these areas could send tactile information directly or relay tactile signals through any one of the higher visual cortical areas projecting to primary visual cortex. Given that we did not find evidence for whisker activation in primary visual cortex neurons, perhaps feedback from RL is selective to the primary modality of the target structure and carries solely visual information to V1. Experiments using axonal imaging may elucidate the functional role of long-range projections from higher visual areas to primary visual cortex.

Cross-modal plasticity following the loss of visual inputs has been shown to give rise of tactile signals in primary visual cortex (Amedi et al., 2010; Merabet and Pascual-Leone, 2010; Van Brussel et al., 2011). Human neuroimaging studies have shown V1 activation in blind subjects during braille reading and other tactile paradigms (Merabet et al., 2008; Sadato et al., 1996). Cross-modal activity in primary visual cortex was also observed after short-term visual deprivation and causally linked to tactile sensation (Merabet et al., 2008, 2007). In rodents, cross-modal plasticity following the loss of visual inputs depends on inputs from the whiskers (Newton et al., 2002; Van Brussel et al., 2011). In this context, our results suggest that cross-modal plasticity in primary visual cortex may require more than the strengthening of existing synaptic connections and involve concerted changes across multiple brain structures.

Overall our results provide strong evidence supporting the idea that primary sensory areas encode mainly their primary sense during naturalistic behaviors and raise questions about the functional role of direct and indirect connections between primary sensory areas.

## AUTHOR CONTRIBUTION

JC, SK and VB designed the experiments. JC and SK performed the experiments. DM and SK designed and performed preliminary experiments. JC analyzed data. JC, SK and VB wrote the paper with input from DM, BLM and LA. VB supervised the project. VB, BLM and LA secured funding.

## ACKNOWLEDGMENTS

We are grateful to Karl Farrow and members of the Bonin and Farrow laboratories for feedback, discussions and comments on the manuscript. We thank Jessica Sternisa and Paula Rodriguez for animal training. We thank Adrian Cheng for technical advice with the two-photon microscopes. This work was supported by Neuro-Electronics Research Flanders (BLM and VB), AIHS Polaris award (BLM), AIHS graduate studentship (DM), NSERC funding (BLM), NSF grant 1631465 (BLM), Research Foundation—Flanders (FWO) grant G0D0516N (VB), and KU Leuven Research Council grant C14/16/048 (VB,LA).

## MATERIAL AND METHODS

### Animals

All animal procedures were approved by the Animal Ethics Committee of KU Leuven. We report on 11 mice. Of these, 3 were C57Bl/6j mice (22 to 30 gr, 2 to 5 months), n = 3 were Thy1-GCaMP6 mice (Dana et al., 2014) and 5 TRE-CaMKII-GCaMP6 (Wekselblatt et al., 2016). Mice were implanted with a head plate and trained to run on a 150-cm linear treadmill belt for a periodic water reward (Mao et al., 2017; Royer et al., 2012). All mice were implanted with a cranial window for chronic cellular imaging above posterior cortex (Goldey et al., 2014).

### Surgical Procedures

Mice were injected with dexamethasone (3.2 mg/kg I.M., 4 h before surgery), anesthetized with isoflurane (induced 3 %, 0.8 L/min O_2_; sustained 1–1.5 %, 0.5 L/min O_2_), and implanted with a titanium head plate. For cellular imaging in V1 and S1, a craniotomy was made, and 5-mm cranial glass window implanted over left posterior cortex (2.5 mm anterior to lambda, 2.5 mm lateral to midline). Head plate and cranial windows were affixed with dental cement (Metabond, Crown & Bridge and Kerr Tab, Kerr Dental) mixed with charcoal to provide light shielding for imaging. All mice received post-operative treatment for 60 hours (buprenorphine 0.2 mg/kg I.M. and cefazolin 15 mg/kg I.M. in 12-hour intervals) and were given five days to recover.

### Viral Vector Injections

Mice were anesthetized as described above and the cranial windows removed. An adeno-associated virus (AAV) construct containing GCaMP6 and the synapsin promotor (AAV1.Syn.GCaMP6m.WPRE.SV40, U Penn Vector Core) (Chen et al., 2013) was injected to monocular V1 (guided by flavoprotein imaging). 500 nL AAV solution were injected at cortical depths of 250 to 450 μm. The AAV solution contained 25 % D-Mannitol (10 % in PBS) to increase transfection efficacy. Injections were performed using beveled glass capillaries (~20 μm tip diameter, Drummond Sci.) at low injection rates (50 or 100 nL/min) using a microliter injection system (Nanoject II, Drummond Sci.). Cranial windows were replaced, and mice allowed to recover as described.

### Treadmill Assay

The treadmill assay was adapted from (Royer et al., 2012). Two 3D-printed 10-cm diameter lightweight treadmill wheels mounted on a custom frame (Thorlabs) held a 150-cm long, 50-mm wide belt made of Velcro (Country Brook). Cues consisted of 0.5 cm wide strips of foam attached to the belt (4 stripes spaced 1cm per cue). In some cases, the treadmill cues were covered from the animals’ eyesight by a shield mounted 10 to 15 mm in front of the animals’ nose and 1 cm above the belt (–45 to 45 deg. azimuth, –30 deg. elevation). Teflon tape (CS Hyde) was adhered to the platform to reduce friction. A rotary encoder (Avago Tech) attached to treadmill shaft was used to monitor treadmill rotation and belt position at a resolution of 3.14 mm. Once per treadmill rotation, for reward delivery, a photoelectric sensor (Omron) detected a reflective strip attached to the underside of the belt triggering opening of an electromagnetic pinch valve (MSscientific) and controlling water delivery through a spout. A custom Arduino based circuit board monitored behavior variables and controlled valve opening. Board design and control software is available at https://bitbucket.org/jpcouto/lineartreadmillrig.

### Behavioral Training

Mice were habituated to handling for three days prior to all procedures. Five days after surgery, water intake was restricted to 1 mL per day and animals were trained to head-fixed treadmill locomotion on belts without tactile stimuli. Mice were rewarded with tap water or 7% sucrose solution at the end of each lap (5-20 μL drop size). No visual stimuli were presented during training. Training duration was increased gradually from a few minutes to 1 hour per day over a period of two weeks. Training was completed when animals reached desired levels of performance (>3 laps/min.).

### Flavoprotein Imaging

Retinotopic mapping with flavoprotein imaging was used to guide viral injections. Light excitation was done with a blue LED (470 nm, Thorlabs) and collection through a green filter (510/84 nm filter, Semrock). Imaging was done at a frame rate of 5 fps using a 2x lens (NA = 0.055, Edmund Optics) and an EMCCD camera (EM-C^2^, QImaging; 1004 by 1002 pixels, 4 by 4 binning). Fractional changes in fluorescence were normalized to baseline and averaged across 4-sec intervals to capture the slow time course of the flavoprotein auto-fluorescence signal. The location of monocular V1 was identified by eye to guide targeted viral vector delivery of the genetically encoded calcium indicator GCaMP6 at retinotopic locations corresponding to monocular V1.

### Two-Photon Imaging

A custom-built volume scanning two-photon microscope (Neurolabware) was used to image somatic calcium signals of S1 and V1 neurons in layer II/III (150 to 300 μm below the pial surface) at frame rates of 30 fps. GCaMP6 was excited at 920 nm using a MaiTai DeepSee laser (Spectra Physics / Newport) through a 16x lens (NA = 0.8, Nikon) and green light emission was collected using a green filter (510/84 nm, Semrock) with a GaAsP photomultiplier tube (Hamamatsu). Maximal laser power output at the objective was 20 to 100 mW, depending on the depth of field-of-view. We used a black imaging chamber and blackout material (Thorlabs) to block stray light from the visual display.

### Visual Stimulation

For visual stimulation, a calibrated 22-inch LCD monitor (Samsung 2233RZ, 1680 by 1050 pixel resolution, 60 Hz refresh rate, average luminance XX cd/m^2^) was positioned 18 cm in front of the right eye, covering 120 by 80 degree in the right visual field (0 to 120 deg. central to peripheral and ±40 deg. lower to upper visual field). Custom software based on Psychopy (Peirce, 2007) was used to control visual stimulation and synchronize the recordings. For the experiment in darkness, the screen was switched off and all light sources covered with blackout material. Light levels were at the detection threshold of our luminance meter (< 0.01 cd/m^2^).

### Eye Tracking

Eye position and pupil size were measured with an infrared camera (AVT Prosilica GC660) and a zoom lens (Navitar Zoom 6000). Infrared light was focused on the eye using an LED light source (850 nm, Thorlabs) and a collimated lens (Thorlabs). Data was acquired at >30 frames per second using custom software (https://bitbucket.org/jpcouto/labcams).

### DATA ANALYSIS

All data were analyzed using custom scripts written in Python or Matlab (Mathworks).

### Calcium Imaging Data

Images were registered to an average composed of 1000 frames from the middle of each session using phase correlation. Regions of interest (ROIs) of active neural cell bodies were identified manually using a pixelwise local spatiotemporal correlation criterion (3 by 3 pixels neighborhood, threshold at correlation coefficients > 0.95)(Smith and Häusser, 2010). Raw calcium time courses were calculated by averaging pixel intensities over each ROI and subtracting an estimate of neuropil contamination. The neuropil signal was computed by averaging a ring of pixels around ROIs and using a low-rank SVD approximation. Raw calcium time courses were expressed as fractional changes above baseline fluorescence (dF/F_0_). Baselines were computed by linear regression to the lowest 10 % of the raw time courses. dF/F_0_ time courses were deconvolved to estimate firing rates.

### Behavioral Data

Rotary encoder increments were used to calculate treadmill position at centimeter precision and instantaneous treadmill speed in cm/s. After every completed lap, encoder increments were reset to zero to prevent potential accumulation of treadmill slip. Camera frames from the eye tracker were smoothed with a Gaussian filter, contrast-threshold, and binarized, resulting in black-and-white images. For every image, eye position and pupil diameter were detected by fitting an ellipsis to the pupil. Pupil size was calculated from the equivalent diameter of the ellipsis and expressed in mm^2^. Eye position was expressed as relative change in degree relative to the average eye center position within individual experiments. Artefacts in the data resulting from e.g. eye blinks were removed using a threshold criterion (mean ± 2-times s.d.). Encoder and eye data were resampled at the frame rate of the two-photon microscope.

### Position-Related Analysis

A standard position-related procedure was used to relate calcium time courses to location on the treadmill. The treadmill lap was divided in 150 1-cm intervals. Average deconvolved calcium activity was computed for each interval for each lap and normalized by the time the animal spent at each interval, resulting in position-related activity profiles. Accordingly, raw calcium time courses and locomotion speed were normalized to treadmill location with the same procedure. Trial-to-trial reliability of activity profiles was measured by computing the fraction of variance in single trials that is explained by the average across laps. Formally, the measure of explained variance follows EV position = (P_r_–P_e_)/P_r_*100, where P_r_ is the variance of the single trial responses and P_e_ is the mean square distance between single trial responses and the across trial. EV was two-fold cross-validated (100-times) estimating how a random half of the trials predicts the other half.

