## Supplementary figures and images for "Spatially segregated responses to visuo-tactile stimuli in mouse neocortex during active sensation"

### Supplementary Figure 1

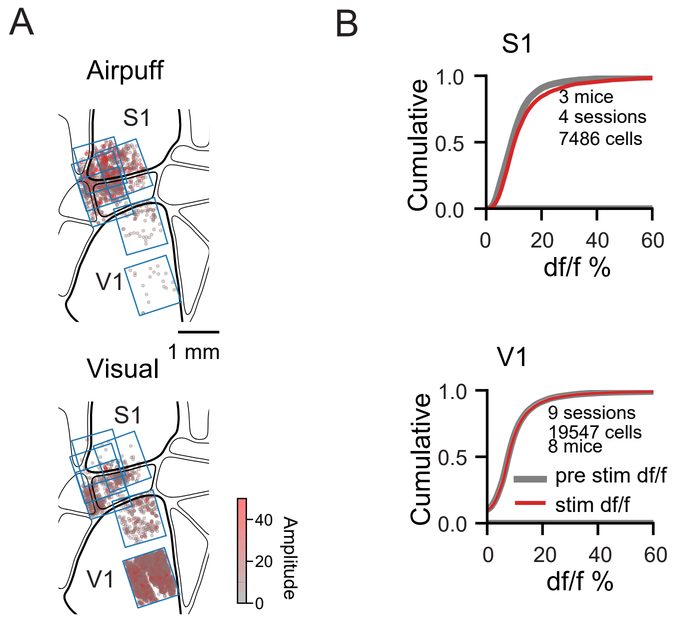

### Supplementary Figure 2

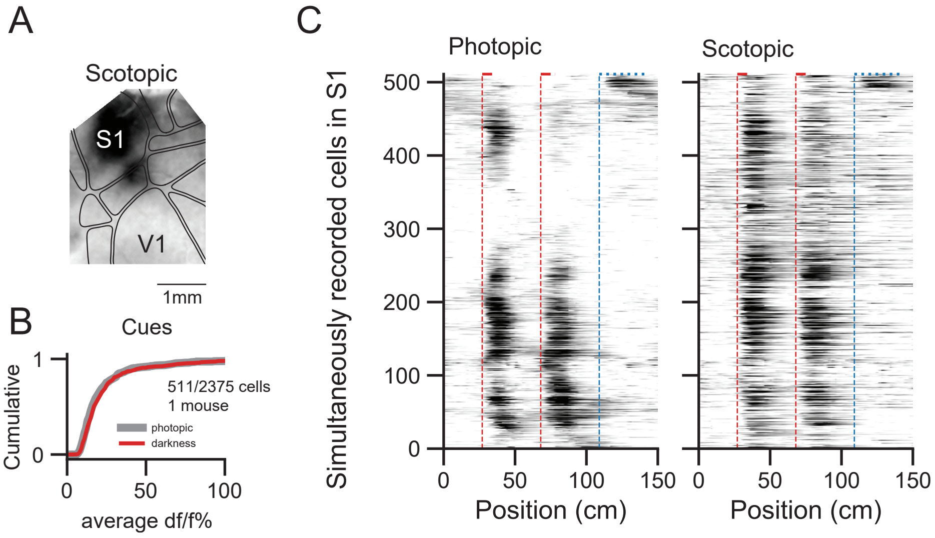

### Supplementary Figure 3

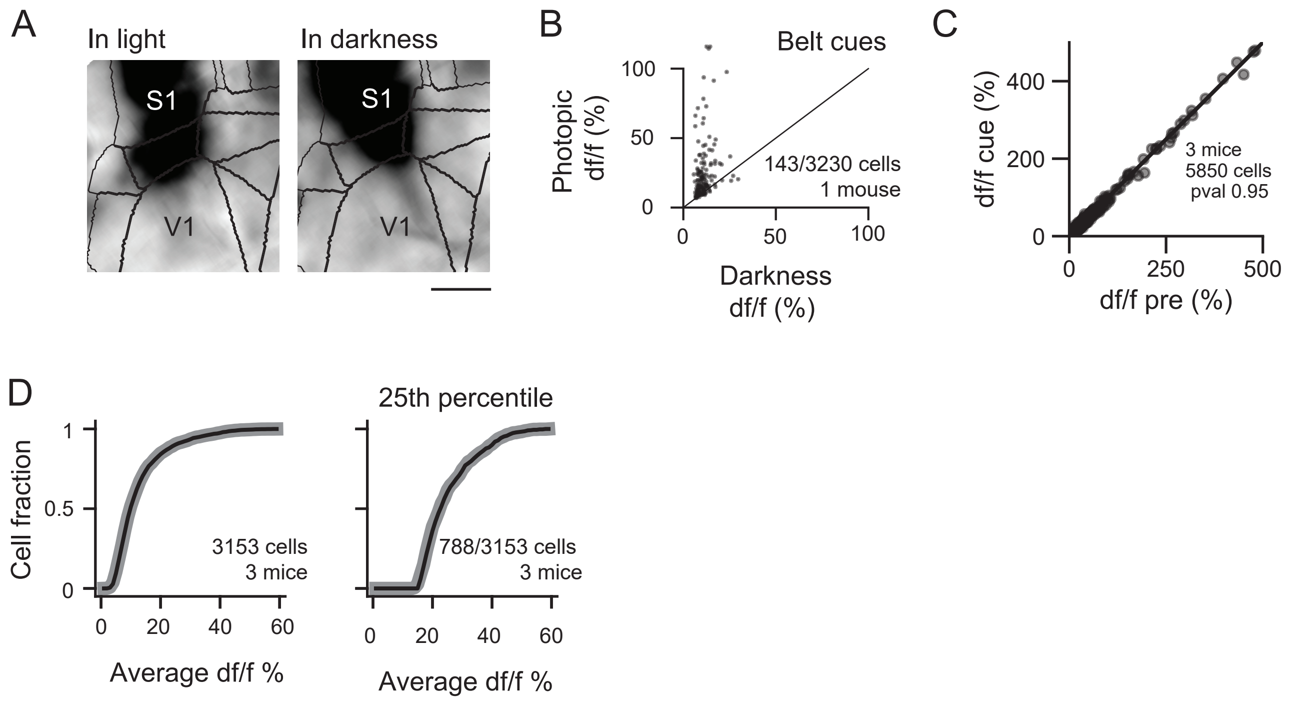
